# Digging deeper into the immunopeptidome with TripleToolWF

**DOI:** 10.64898/2026.06.11.731513

**Authors:** Rupert L Mayer, Karl Mechtler

## Abstract

While the field of immunopeptidomics has matured substantially over the last years, high input amounts of cellular or tissue material are still required to obtain a somewhat complete profile of the immunopeptidome. Here we present a simple platform termed TripleToolWF (derived from Triple Tool workflow) to increase the number of identified and quantified immunopeptides combining the outputs of three search engines such as PEAKS Online 12, Sequest HT with INFERYS rescoring and MSFragger. For assessing the false discovery rate (FDR) an entrapment approach is used. The platform improved peptide identifications by 6-14% and peptide quantitations by 11-25% compared to the best individual search engine for two independent, previously published, bacterial infection datasets. Peptides were mostly 9-12mers as expected and >90% of the obtained 9mers were predicted binders by the stringent majority voting approach of Immunolyser 2.0 which indicates high confidence of the identified immunopeptides. The FDR was monitored using dedicated entrapment searches against shuffled databases. The resulting entrapment FDR was assessed before and after result pooling and showed only a minor increase upon pooling compared to the worst individual search engine. It remained even below the target of 1% peptide FDR in 40% of the experiments. Compared to the original publications, the number of high confidence bacterial immunopeptides was drastically elevated by 53% and 2800% for the Listeria monocytogenes and Mycobacterium bovis BCG projects, respectively, when applying strict filters. Of these additional bacterial sequences, >92% of the 9mer sequences were predicted as binders by at least one of the prediction algorithms of Immunolyser 2.0 illustrating their actual HLA binding nature. TripleToolWF hence provides a simple tool to further increase the number of obtained sequences from MS-based immunopeptidomics experiments to facilitate a deeper view of the immunopeptidome for refined vaccine candidate prioritization.

## INTRODUCTION

Immunopeptidomics has evolved as an important complementary technology for vaccine development as MHC-presented epitopes can directly be detected. Sensitivity in immunopeptidomics has experienced quantum leaps since the initial studies in the Rammensee and Hunt labs.^1,2^ While only around a hand full of peptides was identified for a particular cell line in the early 90s, nowadays typically thousands of peptides can be confidently identified by modern instrumentation and software tools from a fraction of the sample material used back then. Compared to tryptic peptides, both the measurement and data analysis of these short peptides is hampered by alternative ionization and fragmentation patterns. This poses challenges for all classical proteomics search engines as they were designed predominantly for tryptic peptides. However, rescoring the initially identified peptide-spectrum matches (PSMs) by utilizing additional features such as fragment intensities, retention time and others can substantially boost identification of immunopeptides, other non-tryptic peptides and even tryptic peptides ^3–6^. Willems et al. have further demonstrated that pooling the results of multiple search engines before rescoring further improves identification rates for immunopeptides for a bacterial infection dataset^7^.

Here, we propose an alternative workflow termed TripleToolWF that combines the immunopeptide identifications of three search engines, namely PEAKS Online 12, Sequest HT with INFERYS rescoring and MSFragger. Rescoring happens at the level of each individual search engine before peptide sequence pooling followed by assessment of the combined false discovery rate (FDR) using entrapment searches. Two immunopeptide studies focusing on bacterial infection were reassessed with TripleToolWF leading to markedly elevated numbers of identified and quantified peptides compared to the original studies. When comparing the pooled results to the best individual search engine, both peptide IDs (+6-14%) and peptide quantitations (+11-25%) were noticeably improved. While the FDR of the combined results was slightly elevated compared to the single search tools, false discovery rates were still very well controlled at <3% and <1% for the Listeria and Mycobacterium BCG study, respectively. The number of high confidence bacterial peptides identified was drastically elevated by 36 (+53%) for Listeria and 28 (+2800%) for BCG. The additional 36 Listeria peptides represent a further improvement to the 18 additional bacterial peptides reported by Willems et al. on the same original data.

## RESULTS

### TripleToolWF uses parallel target and entrapment searches to control peptide pool FDR

All raw data deposited on at the PRIDE repository from Mayer et al. 2021^8^ (PXD031451, referred to as “Listeria study” going further) and Bettencourt et al. 2020^9^ (PXD015646, referred to as “BCG study” from now on) were downloaded and searched using PEAKS Online 12^10^, Proteome Discoverer 3.2 together with Sequest HT and INFERYS rescoring^11^ as well as MSFragger^12^. Detailed settings applied for the searches are provided in the materials and methods section. Identified peptide sequences were pooled, duplicates removed, and only unique stripped peptide sequences were retained. In parallel to the target database searches, an entrapment database search was carried out for each software to assess the FDR before and after peptide ID pooling. Pooling of peptide sequences could lead to substantially elevated numbers of false positive sequences in the final list due to accumulation of software-specific false positives but reproduction of true positive peptide sequences. Therefore, it is imperative to monitor the FDR before and after peptide sequence pooling. In case of unsatisfactory FDR control, stricter target FDRs can be adopted both for the target and the entrapment searches as indicated in the workflow overview (Fig. 1). In case of acceptable FDR control, the target ID list of the target database search is further filtered for quantified peptides only and peptide length and binding assessed to further assess the quality of the obtained immunopeptide sequences.

**Figure 1:**
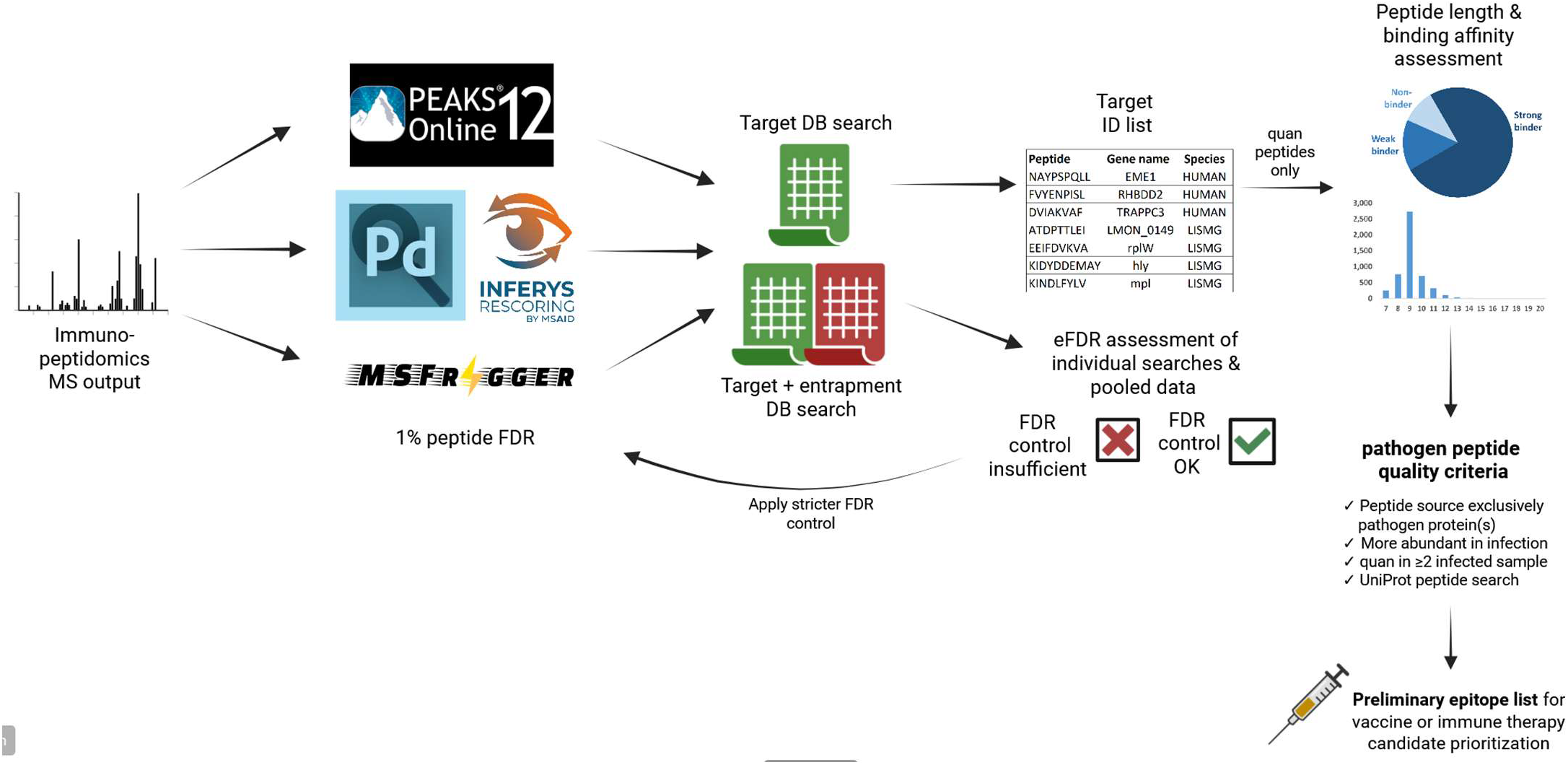
Schematic overview of the TripleToolWF combining the results of the three different tools PEAKS Online 12, INFERYS rescoring and MSFragger to improve immunopeptide ID coverage. The potentially elevated FDR from combining the results is tightly monitored using an entrapment approach allowing application of stricter FDR settings to achieve the desired FDR.

### Reanalysis by TripleToolWF boosts MHC I peptide identifications and quantitations

The Listeria study consisted of samples from two different cell lines (HeLa and HCT-116) either infected with Listeria or uninfected. These were measured by LC-MS both as label-free and TMT-labelled peptides with four replicates per condition resulting in a total of 32 LC-MS runs. For the BCG study, only data from one of the four experiments (experiment 4) was publicly available and therefore utilized in the current manuscript. These data included MHC class I samples derived from a single cell line (THP-1) with two different uninfected conditions (+/-cytokines), and four infected conditions (+/-cytokines, live or heat killed BCG) with two fractions per condition and two replicates per condition resulting in a total of 24 LC-MS runs. While exact numbers of all pooled peptide IDs are not reported for both studies, the number of pooled, stripped peptide sequences per study is estimated to a total of around 18,000 and 16,500 immunopeptides for the entire Listeria study and experiment 4 of the BCG study, respectively. While these numbers represent already a respectable immunopeptide coverage, the TripleToolWF was able to further extend the depth of analysis to 20,910 and 20,968 peptide sequences for the Listeria and BCG study, respectively (Fig. 2A, Supplementary Data 1-4). With 18,386 and 19,886 peptides for the Listeria and BCG study, respectively, PEAKS contributed most IDs to the pool, with additional contributions from MSFragger and INFERYS. Beyond peptide identifications, elevated numbers of peptides were also quantified with +11% for the Listeria and +25% for the BCG study. While peptide quantification is underutilized in many immunopeptidomics studies, it can severely strengthen the confidence of disease-specific immunopeptides particularly when differential abundance is considered between control and diseased samples ^7,13^. Peptides were considers successfully quantified when at least two quantified values were reported in at least one condition. For the Listeria project this means that at least in 50% of the replicates of at least one condition a peptide was quantified. For the BCG project this translates to two samples of the same biological replicate when technical replicates and fractions are treated as replicates.

**Figure 2:**
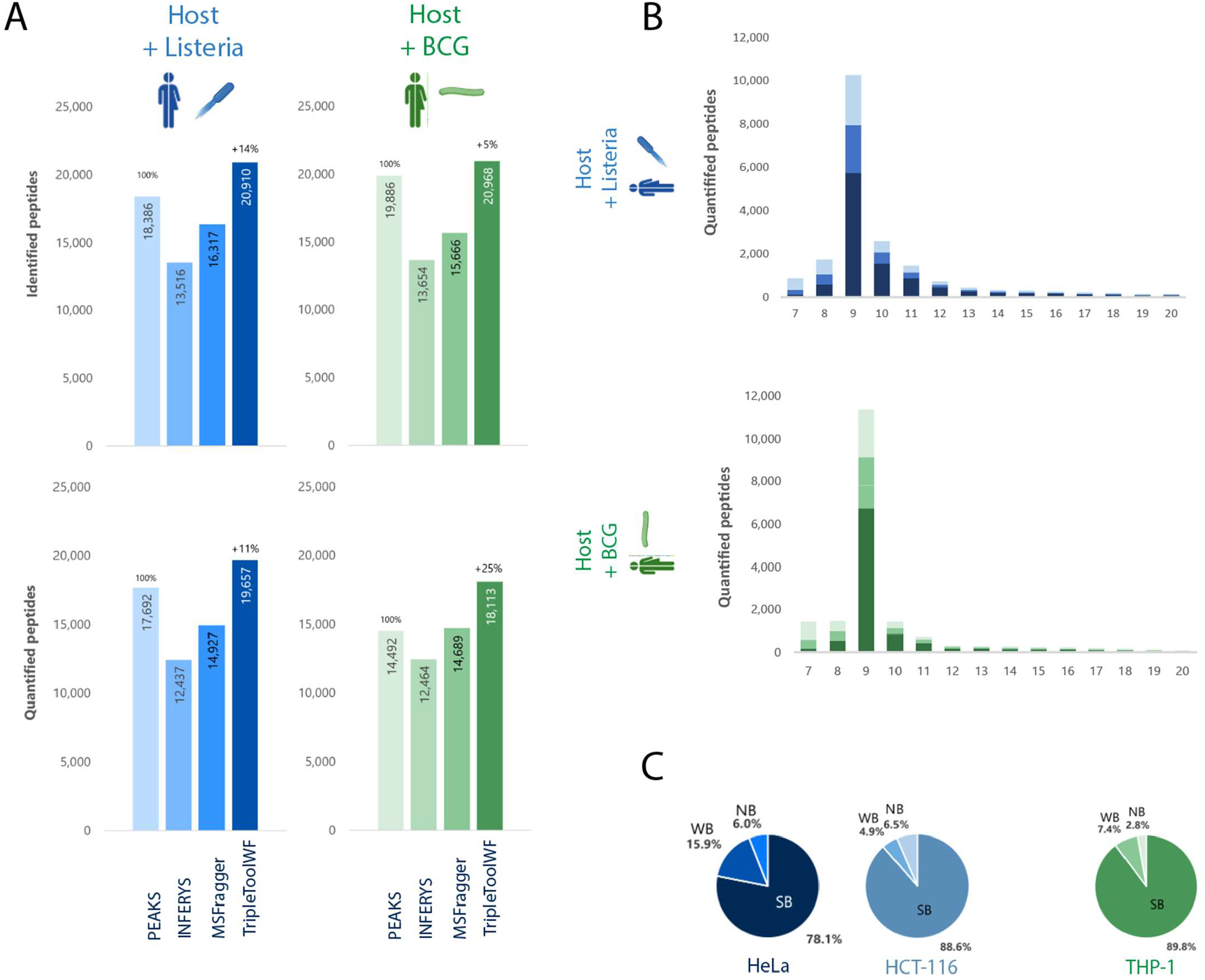
TripleToolWF results in elevated number of peptide identifications and quantifications for host and bacterial immunopeptides. **A** The TripleToolWF was tested on two bacterial infection data sets showing increased number of quantified peptides compared to the original studies. Combining the results of the three tools increased peptide IDs by 5-14% and quantified peptides by 11-25% compared to PEAKS Online 12 alone. **B** Peptide length distribution for quantified peptides. Peptides quantified by all three tools are shown in the darkest color, followed by peptides quantified by two tools (medium brightness bars) or a single tool (brightest color). **C** Identified 9mer peptides were submitted to Immunolyser 2.0 majority voting using binding predictors NetMHCpan 4.2, MHCflurry 2.0 and MHCpred 3.0. Venn diagrams indicate fractions of strong binders (SB), weak binders (WB) and non-binders (NB).

The quantified peptide sequences follow the expected peptide length distribution of mostly 8-12mers (Fig. 2B, Supplementary Data 5) very well and most of the quantified peptides are quantified by all three software tools (darkest bar section) followed by peptides quantified by two tools and the lowest contribution coming from uniquely quantified peptides. Immunolyser 2.0 was utilized to assess the binding affinity resulting in a high proportion of binders among the identified 9mer sequences (Fig. 2C, Supplementary Data 6 & 7).^14^ Strong and weak binders contributed at least 93.5% of all identified 9mer peptides. This further substantiates the relevance of the identified peptide sequences.

### Many additional bacterial pathogen peptides are identified

TripleToolWF also demonstrates a substantial increase on the identification and quantification of additional bacterial peptides, (Fig. 3). This is of particular importance due to the high translational relevance of these sequences. To qualify peptide sequences as high confidence bacterial peptides, quantified bacterial peptides were filtered for i) increased abundance upon infection, ii) at least two quantifications in at least one infected sample condition, and iii) no I/L permutated sequences in human upon UniProt peptide search. Indeed, 36 additional Listeria-derived (+53%) and 28 additional BCG-derived (+2,800%) peptides were reported as high confidence bacterial immunopeptides (Fig. 3A). Nearly all Listeria peptides from the original study were recovered, with only four peptides lost. Most of these lost peptides, however, are predicted as non-binders and are therefore potentially false positives or of only minor translational interest. For the BCG study, the presented workflow recovered the single peptide of the initial study that met the applied criteria employed in the reported work. The additional Listeria peptides fulfill the expected length distribution of mostly 8-12mers while the additional BCG peptides feature a slightly elevated fraction of 7mer peptides (Fig. 3B). 7mers are typically a minor fraction of peptides eluted from MHC class I molecules. However a few examples are reported to bind selected MHC class I alleles ^15^ and evoke a CD8^+^ immune response.^16,17^ Binding prediction of the TripleToolWF-specific bacterial immunopeptides using Immunolyser 2.0 indicates 85.2% majority voted binders for Listeria 9mers, but only 53.9% majority voted binders for BCG 9mers (Fig. 3C). When assessing the percentage of 9mer peptides predicted as binders by at least a single predictor, this figure increases to 100% and 92.3% binders for Listeria and BCG, respectively (Fig. 3D). While for Listeria the discrepancy between majority voted binder and single tool predicted binder is rather small (~15%), BCG shows a remarkable 38.4% difference. This is partially explained by the HLA alleles present in the different studies: while the Listeria study employed HeLa and HCT-116, with a total of ten different MHC class I alleles, the BCG study featured only five alleles, of which two are not covered by the most promiscuous prediction algorithm MixMHCpred. While for the Listeria study only for a single allele MixMHCpred prediction was not possible. MixMHCpred is typically the least stringent of the three binding predictors and returns the highest number of predicted binders. Therefore, the incompatibility of these two alleles (HLA-B*15:15, HLA-C*03:13) with MixMHCpred prediction at least partially explains the stark discrepancy between the fraction of majority voted binders and single tool predicted binders.

**Figure 3:**
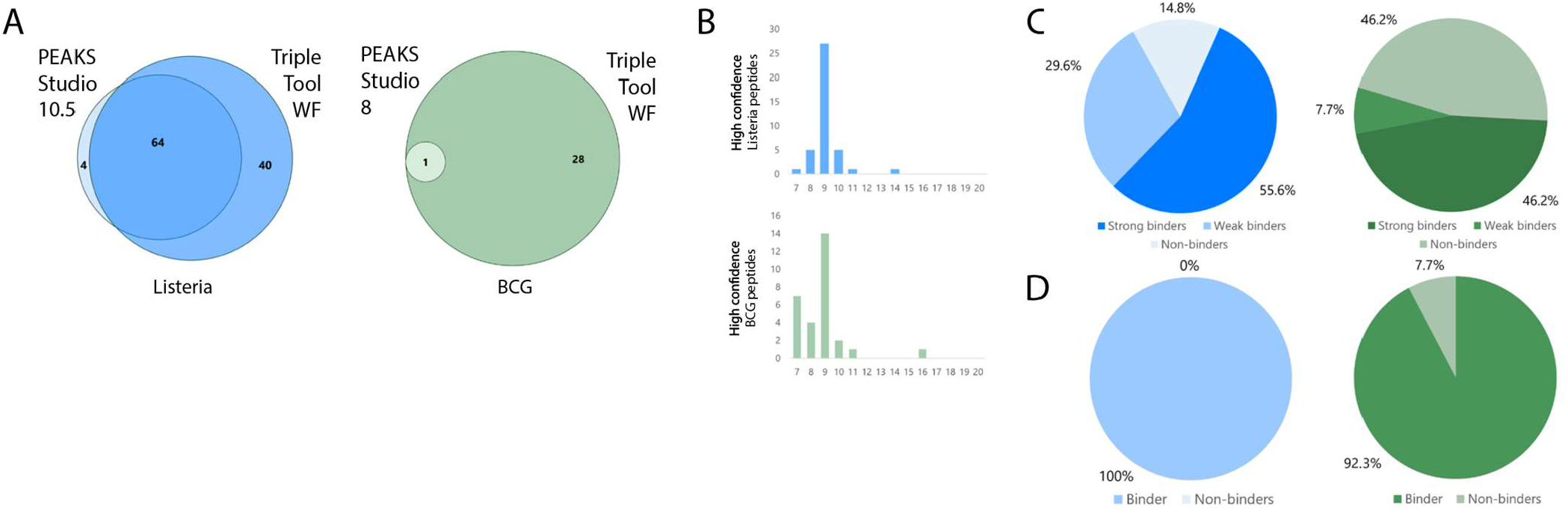
Elevated numbers of high confidence bacterial peptides. **A** Quantified pathogen peptides for the TripleToolWF were filtered for i) higher abundance upon infection, ii) at least two quantifications in at least one infected sample condition and iii) no I/L permutated sequences identified in human upon UniProt peptide search. TripleToolWF reported +53-2800% more high-confidence bacterial peptides increasing bacterial epitope depth. For both, Listeria (left Venn diagram, blue) and BCG (right Venn diagram, green), considerable overlap of the reported high confidence bacterial peptides with the original studies is demonstrated. While four Listeria peptides were lost, no BCG peptide was lost. **B** Peptide length distribution mostly matches expected dominance of 8-12mers. **C-D** High confidence novel bacterial 9mers were subjected to binding prediction using Immunolyser 2.0 i) majority voted binding (**C**) and ii) binding prediction by at least a single tool (**D**).

### Pooling identifications results in only minor increases in false discovery rates

Since TripleToolWF involves pooling results from three different software tools, it is imperative to assess and properly control the false discovery rate. The entrapment databases were generated by dbtoolkit-4.2.5 using the shuffle databases option^18^. Raw data were searched against human + respective bacterial + contaminant database + all of the three databases shuffled. Target FDR was set to 1% on all levels, only protein FDR was set to 100%. Peptides uniquely identified from entrapment DB proteins were counted as entrapment hit. Sequences matching both target and entrapment proteins were excluded. The resulting global entrapment FDR (eFDR) is slightly elevated, but still remains quite well controled (Fig. 4). This holds particularly true when compared to the respective software tool with the highest eFDR. The highest eFDR is observed mostly for PEAKS Online 12 which is potentially resulting from the the unique decoy fusion method that PEAKS employs ^15^. For the Listeria study, the eFDR is interestingly elevated to around 3% for the label-free samples. However, TMT-labeled data is well below 2% and even below 1% for the HCT-116 TMT data. For the BCG study, the eFDR for the TripleToolWF still remains substantially below 1% at 0.63%. Taken together these results demonstrate adequate FDR control for the presented approach for these two datasets.

**Figure 4:**
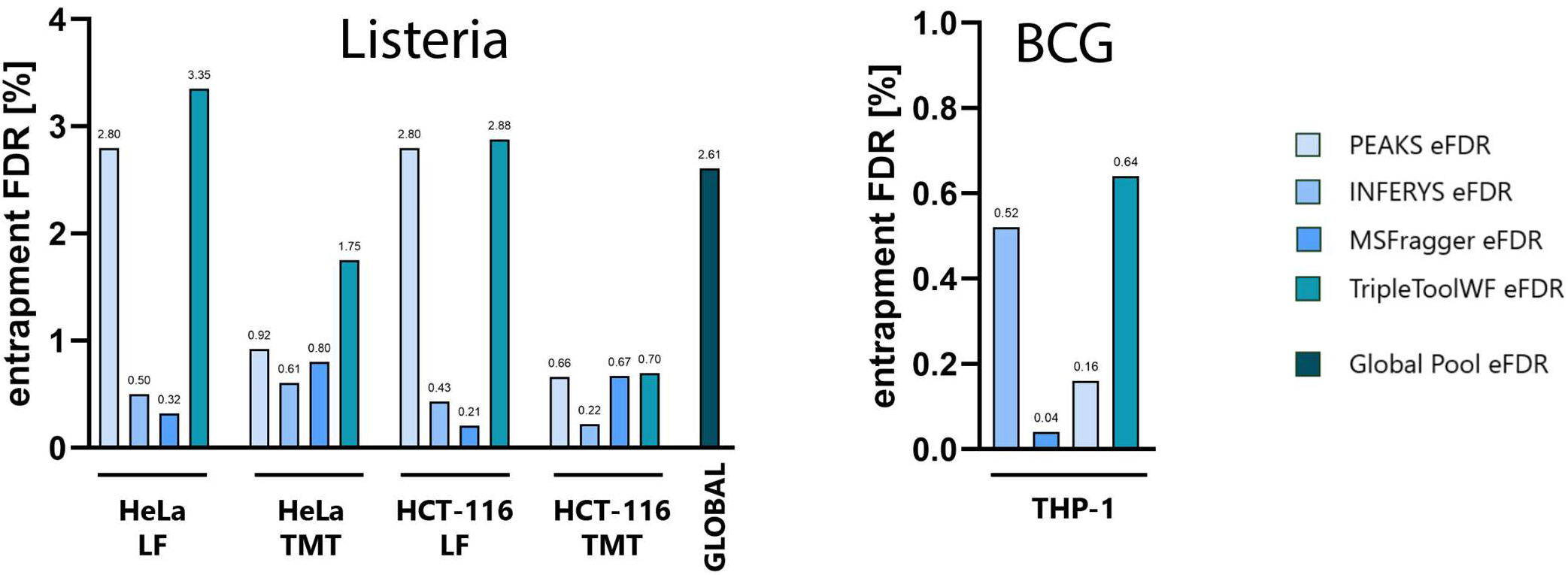
eFDR assessment of single and pooled searches. Raw files were searched against a combined database of target and shuffled human + bacteria + contaminant fasta files to assess the FDR increase peptide sequence pooling. While the FDR is increased after result pooling, the overall FDR remains well controlled and only slightly elevated compared to the highest FDR single tool analysis. Only for the Listeria HeLa TMT samples the resulting combined eFDR was markedly increased with +0.83% eFDR. The target FDR in all cases was set to 1% on PSM and peptide level (100% FDR on protein level).

## DISCUSSION

Despite substantial improvements in sample preparation, LC-MS analysis and data analysis, the field of immunopeptidomics still suffers from incomplete analysis depth as compared to other subdomains of proteomics like single cell proteomics. This is mainly caused by the ultra-low abundance and non-tryptic nature of the MHC ligands. In order to circumnavigate these challenges, innovative strategies during sample preparation, measurement and data analysis are needed. TripleToolWF, the presented tool in this manuscript, describes a simple methodology to improve the immunopeptidomics data analysis for a deeper picture of the immunopeptidome. This method utilizes simple pooling of identified stripped peptide sequences from three different software tools with an added entrapment strategy to monitor FDR control. The strategy increases the overall number of identified peptides over each individual software output by up to 14% and the number of quantified peptides by up to 25% as assessed by reanalyzing two distinct bacterial infection datasets. The pooled immunopeptide sequences match the expected peptide length distribution well and >90% of the quantified 9mers are predicted binders, which indicates high confidence of the immunopeptidomics nature of the obtained sequences. Concerning the peptide FDR after the result pooling, only a minor elevation over the worst individual tools FDR was observed for the large majority of the samples. This indicated that the FDR is still controlled well resulting in a peptide entrapment FDR of below 1% for two of the five experiments. Further filtering the pooled data for high confidence bacterial peptides achieved a considerable gain of 36 and 28 additional peptides for Listeria and BCG, respectively. These additional peptides featured mostly 9mers as expected but also an unexpectedly high fraction of shorter peptides for BCG. 7mer peptides are a rather minor fraction of immunopeptides in general but presentation of immunologically relevant examples has been reported.^16,17^ While many of the high confidence 9mer bacterial peptides are predicted as binders using the Immunolyser 2.0 majority voting, particularly the novel BCG peptides demonstrate mostly non-binders using the strict majority voting option. However, when lowering the stringency to require only a single algorithm to predict as binder, all 9mers from both Listeria and BCG are predicted binders indicating relevant binding affinity of the additional bacterial peptides. The novel Listeria sequences included peptides from well-known virulence factors hly, mpl, plcB, which had also been identified in the original work, as well as other proteins for which peptides had been identified in the original work (PdhD, prsA, rplW). The additional BCG sequences included three peptides (FSRPGLPV from FBPA, FBPB or FBPC, TLHEVPVL from BCG_3864C, WDCAAVNV from BCG_1242C) that were previously reported as epitopes from Mycobacterium according to the Immune Epitope Database (IEDB, https://www.iedb.org/) ^19^. Furthermore, 12 other novel sequences originate from proteins that are also reported in the IEDB as Mycobacterium antigens including NUOJ, DNAK, FAR, TUF, BCG_3789, BCG_0261C, METC, PE_PGRS11, SUGI, TOPA, FADE35 and SPPA. This substantial overlap with previously reported MHC-presented sequences further corroborates the significance of the obtained high confidence bacterial epitopes.

Willems et al. have previously described an approach that similarly leverages multiple search engines including PEAKS, Comet, Sage and MSFragger to boost immunopeptide identification rate.^7^ Their workflow further utilizes rescoring with MS^2^rescore of each software output before integration into a single output file using mokapot for data integration and peptide q value control at ≤0.01.^20,21^ Willems and colleagues demonstrate a substantial improvement with 18 high confidence Listeria peptides more than the original work and increase the confidence from 5% to 1% peptide FDR, simultaneously increasing both analysis depth and confidence. TripleToolWF, however, further extends the number of identified high confidence peptides even though the FDR is slightly less controlled than for the work of Willems et al.. Quantification in the workflow of Willems et al. is performed using FlashLFQ or IsobaricAnalyzer.^22^ One minor pitfall of their workflow, however, is that FlashLFQ does not support quantification of Bruker timsTOF data (at the date this manuscript was compiled). Contrastingly, quantification for TripleToolWF is achieved by each respective software tool individually with PEAKS Online 12 and MSFragger capable of quantifying Bruker timsTOF data, while PD3.2 employing Sequest HT and INFERYS is not compatible with quantification in Bruker timsTOF data. As INFERYS provided the lowest number of additional peptide sequences, this can be neglected when analyzing Bruker timsTOF data or another search engine could be utilized such as SAGE or Comet.

## METHODS

### Reuse of Listeria study data

All raw files and fasta files from the original publication “Immunopeptidomics-based design of mRNA vaccine formulations against Listeria monocytogenes” have been downloaded from the PRIDE partner repository with the dataset identifier PXD031451.^8,23^ The raw data downloaded from PRIDE represent the complete study.

### Reuse of BCG study data

All MHC class I raw files available from the original publication “Identification of antigens presented by MHC for vaccines against tuberculosis” have been downloaded from the PRIDE partner repository with the dataset identifier PXD015646.^9,23^ These data represent experiment 4 only. Data from experiment 1-3 of the BCG study as well as fasta files were not available for reanalysis.

### PEAKS Online 12 searches settings

Label-free raw data were submitted to the DeepNovo Peptidome Q workflow and “Mass correction” and “Associate features with Chimera Scan” were enabled. MS1 mass tolerance was set to 10 ppm and MS2 tolerance to 0.02 Da. “None” was chosen as enzyme and methionine oxidation and protein N-terminal acetylation selected as variable modifications. Peptide lengths were restricted to 7-20. Protein databases for the respective projects were selected. 1% Peptide FDR was chosen as report filter with 2% confident amino acid threshold and 95% as deep novo score threshold. Deep Novo Protein Association Tag Sharing was set to 5. Label-free quantification was used applying Match between runs with a mass error tolerance of 20ppm and a retention time shift tolerance of 2 min with “Auto Detect” enabled. Feature intensity threshold was set to ≥1,000. Peptide feature thresholds were set to 0 for all, only the charge state was set to 1 ≤ charge ≤ 5. No normalization was applied.

TMT-labeled raw data were submitted to the PEAKS Q workflow, applying mostly the same settings as for the label-free data. Deep learning was enabled and TMT10plex chosen as fixed modification. Again 1% peptide FDR was chosen and no filtering on protein level was performed. “TMT/iTRAQ Label” was chosen as quantification type and TMT-10plex (CID/HCD) with a mass error tolerance of 15 ppm was selected using MS2 as reporter ion type. No purity correction was done and no spectrum filtering performed. Protein filter was set to significance ≥0. No normalization was carried out.

### FragPipe and MSFragger search settings

FragPipe v23.1 was utilized as framework for MSFragger 4.3, IonQuant 1.11.11 as well as DIA-NN 1.8.2 beta 8. The “Nonspecific-HLA” workflow was loaded as basis for the search and was modified so that only methionine oxidation and protein N-terminal acetylation were kept for both TMT and label-free data, and TMT10plex at lysines and peptide N-termini was set as fixed for TMT samples. MSBooster for rescoring was enable using the default DIA-NN model. Quantification was performed using default setting in Quant (MS1) for label-free data and Quant (Isobaric) for TMT data.

### Proteome Discoverer 3.2, Sequest HT search and INFERYS rescoring

For the both the label-free and the TMT searches the processing step consists of the “Spectrum Files”, “Spectrum Selector”, “Sequest HT”, “INFERYS Rescoring” and “Percolator” nodes. In addition, the “Minora Feature Detector” and the “Reporter Ions Quantifier” were specific to the label-free and the TMT database searches.

Analysis templates for both label-free and TMT searches are provided including all workflow details. In brief, spectrum selector was set to include charge 1-6 keeping other settings at default. Sequest HT was set to 10 ppm for MS1 and 0.02 Da for MS2 mass tolerance with a maximum of 2 missed cleavages and no-enzyme (unspecific). Peptide length was specified as 7-20 and dynamic modifications to include methionine oxidation and protein N-terminal acetylation with no static modifications. Percolator was kept at default settings.

The consensus workflow for both label-free and TMT searches included the nodes “MSF Files”, “PSM Grouper”, “Peptide Validator”, “Peptide and Protein Filter”, “Protein Marker”, “Protein scorer”, “Protein Grouping”, “Protein FDR Validator”, “Peptide in Protein Annotation” as well as the post-processing nodes “Result Statistics”, “Data Distributions”, and “Statistics Insights”. “Feature Mapper” and “Precursor Ions Quantifier” were specific to the label-free searches, while “Reporter Ions Quantifier” was specific for the TMT searches.

Peptide and Protein Filter was set to peptide confidence at least low and keeping lower confident PSMs. “Remove peptides without protein reference” was set to false. Minimum number of peptides was set to 1, only count rank 1 Peptides set to False and Count Peptides Only for Top scored Proteins also set to False. All low and medium confidence peptides were filtered out after exporting manually.

### Peptide binding prediction using Immunolyser 2.0

Amongst other functionalities, Immunolyser 2.0 allows assessment of peptide binding using NetMHCpan-4.2 ^24^, MHCpred 3.0 ^25^, and MHCflurry 2.0 ^26^ for MHC class I peptides of 8-14mer peptides. It reports whether a peptide was predicted as binder by majority voting of the different predictors. (“majority voted binders”) but also indicates the individual algorithms predictions. Peptide sequences (8-14mers) were submitted to the Immunolyser 2.0 platform according to the instructions on the webpage (https://immunolyser.erc.monash.edu/). Preferred length of motifs was set to 9 and the alleles for the respective cell line chosen and the default list was chosen for the MHC I typing prediction. The binding predictions for the individual HLA alleles were downloaded and combined.

### FASTA files, databases management & generation of entrapment databases

FASTA files for the Listeria project were reapplied as utilized by Mayer et al. in the original manuscript combining a human + Listeria database (23,260 total combined protein entries) as well as using the MaxQuant v 1.6.3.4 contaminant database (246 protein entries). Since no FASTA files were provided on PRIDE for the BCG project, a more current Mycobacterium bovis (strain BCG / Pasteur 1173P2) fasta file (3,892 protein entries, Dec 2024) was combined with a human uniprot reference fasta file (20,675 protein entries, June 2025) for the BCG project. As no contaminant database was utilized for the original study, none was also applied in the current manuscript. The respective databases (human + Listeria + contaminants and human + BCG) for the target searches were combined using dbtoolkit-4.2.5.^18^ For the generation of the entrapment databases, the combined databases shuffled using the “Output as shuffled FASTA file” option. Both the shuffled and the target databases were then concatenated also using dbtoolkit-4.2.5. All fasta files are provided as supplementary files.

## Supporting information

FileS1 - Listeria - raw exports

FileS2 - BCG - raw exports

FileS3 - Listeria - IDs & quant - processed

FileS4 - BCG - IDs & quant - processed

FileS5 - Lengths

FileS6 - PepBdgAll Listeria Immunolyser

FileS7 - PepBdg BCG Immunolyser

FileS8 - Bacterial peptides - IDs, lengths, bdg aff

FileS9 - Entrapment - Listeria

FileS10 - Entrapment - BCG

FileS11 - Listeria project fasta files

FileS12 - BCG project fasta files

## DATA AVAILABILITY

No new raw data was generated for this manuscript. The original raw data for both studies are available from the PRIDE partner repository with the dataset identifier PXD031451 for the Listeria study and PXD015646 for the BCG study. All data generated during the reanalysis is available as supplementary files.

## ACKNOWLEDGMENTS

The authors are thankful to Nicola Ternette and Paulo Bettencourt for providing additional sample information for the BCG study. The authors furthermore appreciate fruitful discussions with Manuel Matzinger, Fränze Müller and Julia Bubis as well as Peter Pichler on the manuscript. This work was supported by the infrastructure funding fourth call 2022/01 (AT-SCP) of the Austrian Research Promotion Agency (FFG), the FWF projects 10.55776/PAT4142423, 10.55776/PAT4800425 and 10.55776/PAT2059025. We thank the IMP, IMBA, and GMI for general funding and access to infrastructure. For open access purposes, the author has applied a CC BY public copyright license to any author accepted manuscript version arising from this submission.

## AUTHOR CONTRIBUTIONS STATEMENT

RM and KM conceptualized the study, RM performed data analysis and wrote the manuscript. KM edited the manuscript.

